# *Mycobacterium tuberculosis* ESX-T7SS impacts the 3D architecture of tuberculous lesion in susceptible mice

**DOI:** 10.1101/2024.06.14.599018

**Authors:** Caroline G.G. Beltran, Jurgen Kriel, Stefan M. Botha, Margaret B. Nolan, Alessandro Ciccarelli, Ben Loos, Maximiliano G. Gutierrez, Gerhard Walzl

**Affiliations:** Department of Biomedical Sciences, Faculty of Medicine and Health Sciences, Stellenbosch University, Cape Town, South Africa; Central Analytical Facilities, Faculty of Medicine and Health Sciences, Stellenbosch University, Cape Town, South Africa; Department of Physiological Sciences, Stellenbosch University, Stellenbosch, South Africa; The Francis Crick Institute, London, United Kingdom

**Keywords:** CLARITY, 3-dimensional, light-sheet fluorescence microscopy, serial block face electron microscopy, Tuberculosis, granuloma

## Abstract

Tuberculosis (TB) is characterized by the formation of heterogenous, immune-rich granulomas present in various forms in the lungs. Both host and pathogen contribute to this heterogeneity however the molecular and cellular drivers of within-host granuloma heterogeneity remain to be fully elucidated. This knowledge gap is due to a lack of experimental approaches that can fully capture the complex dynamics of the lung architecture, dynamics of host-pathogen interplay and pathogenesis. Here, we developed an approach that combines PACT-based clearing with light sheet fluorescent microscopy for visualizing lesion architecture development and lung involvement in *M. tuberculosis*-infected C3HeB/FeJ susceptible mice. This 3D modelling of whole lung lobes approach revealed critical architectural features in lesion development and lung involvement that were not apparent using traditional thin section imaging. Wild type *M. tuberculosis* infection triggered a clear and well-organized granulomatous-like organization with substantial growth throughout the infection period that invaded a high percentage of the total lung volume. In contrast, infection with the avirulent ESX-1 deletion mutant strain *Mtb* ΔRD1 showed an altered growth pattern with diffuse and sparsely organized CD11b recruitment to sites of infection. Moreover, most of the *Mtb* ΔRD1-triggered lesions were present in the periphery of the lungs and did not display any formal organization. We further provide a novel way of interrogating PACT-cleared tissue for high ultrastructural content using volumetric correlative light and electron microscopy, allowing individual immune cell populations to be tracked and their fate within the granuloma captured. Ultimately, the combination of both modalities allowed an unprecedented view of the architectural distribution of *M. tuberculosis* in the lungs and the progression of lesion development over time. Our data highlight that ESX-1 from *M. tuberculosis* is required for lesion architecture progression in a susceptible mouse model of TB.

## Introduction

Granulomas are complex, multicellular and pathological immune responses localized in the lungs of patients with tuberculosis (TB), representing the hallmark of this disease. It is well established that infection with the bacterial pathogen *Mycobacterium tuberculosis (Mtb)* results in a wide range of pathological heterogeneity ^1,2^, even within the same patient, and that lesion-to-lesion variation determines different outcomes of infection control ^3^. The host pathogen interface plays a dominant role in TB and determines the severity of lung damage ^4^. Patients can present with cavitation, fibrosis or nodular infiltrates, often with a mixture of these pathologies ^5^. It remains challenging to comprehensively characterise this variability and fully understand the implications on the ability of the host to contain the infection and return to normal tissue structure and function. Assessing new drugs, host directed therapies (HDT) and vaccines is still reliant on animal models, more notable murine models, yet suffer from a lack of contextual information specifically regarding molecular and cellular drivers of lesion heterogeneity and the degree and severity of lung involvement. Most studies utilize 2D thin sectioning and/or whole-organ homogenized tissue assessment, providing limited contextual information. Although classic and fluorescent microscopy can be used to observe local host-pathogen interactions in the infected organs, these methods limit the analysis to small areas. Consequently, histologic specimens are subject to fragmented information and sampling bias, and the overall 3D anatomical context is not considered. 3D spatial analysis of infected tissues could allow quantitative profiling while maintaining tissue architecture and identify key biological patterns that facilitates the identification and profiling of important clinical features during TB.

The surge in tissue clarification techniques has unveiled novel ways of interrogating tissue architecture in various biological tissues in three-dimensional (3D) space ^6–9^. These techniques are particularly attractive to study diseases where spatial heterogeneity is typical of the disease. Developments in tissue clarification and deep tissue imaging using light sheet fluorescent microscopy (LSFM) are paving the way for more appropriate ways to visualize tissue without the loss of contextual information ^10–13^. Aqueous-based clearing processes like passive CLARITY (PACT) enables the flexibility of immunofluorescent labelling, whilst retaining endogenous fluorescence to image intact tissues ^8^. PACT-based clearing has been used extensively to study the intact structure of the brain ^7,14,15^, yet its use for lung imaging during *Mtb* infection has been limited ^16,17^. Being able to visualize lesion formation, distribution, and severity of involvement in the lungs, or other organs like lymph nodes, in 3D would greatly enhance our understanding of lesion heterogeneity in the host, enabling us to assess how these develop during TB disease and understand how these resolve with treatment. Coupling this with *Mtb* location and structural organization of different host cell types in lesions like granulomas, will provide a powerful approach to comprehensively characterizing TB disease pathophysiology. The C3HeB/FeJ mouse strain offers the unique potential to model, to a certain extent, the heterogeneity seen in human pathology, with three distinct lesion types visible during low dose infection ^18–20^.

Here, we developed PACT-based clearing and LSFM approach to investigate and map the formation of lesions in 3D in TB-infected C3HeB/FeJ mouse lungs over time. We investigated the dynamics of an ESX-1 deletion *Mtb* mutant and its influence during early and late lesion formation. Following identification of regions of interest in 3D, lesions were excised and prepared for serial block face electron microscopy (SBF) to provide high ultrastructural content from clarified tissue, termed volumetric correlative light electron microscopy (vCLEM).

## Results

### Bacterial burden and lung pathology are reduced in C3HeB/FeJ mice following aerosol infection *Mtb* H37Rv ΔRD1

We used a murine model of TB that more closely represents human pathology by employing the C3HeB/FeJ mouse model and a low dose aerosol infection model using E2 crimson fluorescently labelled *Mtb* H37Rv wild type (E2Crimson *Mtb* WT) and E2 crimson fluorescently labelled *Mtb* H37Rv ESX-1 deletion mutant (E2Crimson *Mtb* ΔRD1) (Fig 1a). Forty mice per group were infected with 100 CFU and the infection allowed to progress over time. At defined time points (1-, 28-, 42-and 70-days post-infection), ten mice per group were euthanized and red blood cells removed via transcardiac perfusion after which lung lobes were processed for passive clarity (PACT), immunolabelling and imaged using LSFM (Fig 1b). PACT based clearing allowed gentle and slow clearing, thereby preserving sufficient lipid content to enable contrast with Electron Microscopy. We therefore identified regions/lesions of interest and further processed these for Serial Block Face Electron Microscopy. Histological analysis and colony forming unit (CFU) determination (Fig 1c) were conducted on the lower right caudal lobes and left lung lobes, respectively.

**Fig 1.**
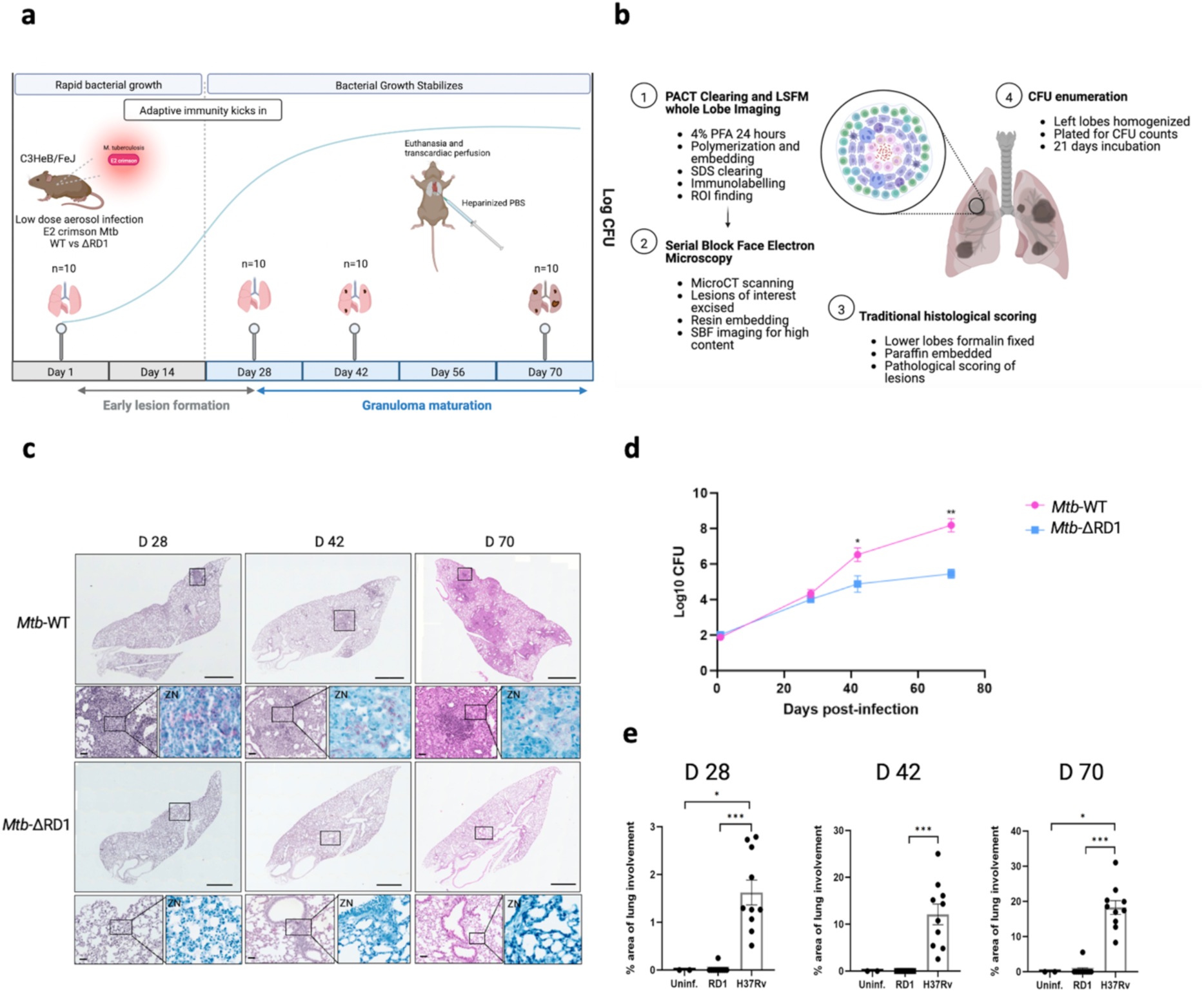
**a** Whole lung lobe imaging workflow. A C3HeB/FeJ murine low dose TB infection model was used. Forty/group of 8–10-week-old C3HeB/FeJ mice were infected via inhalation exposure system with <100 colony forming units (CFU) of either E2Crimson *Mtb* WT (*Mtb*-WT) or E2Crimson *Mtb* ESX-1 deletion mutant (*Mtb-*ΔRD1) *M. tuberculosis* transformed with a PTEC-19 plasmid containing an E2-Crimson reporter. Ten mice/group were euthanized one day post-infection to confirm delivery dose. Infection was allowed to progress over time. Lungs were extracted at day 28, 42 and 70 post-infection (10 mice/group) following transcardiac perfusion with heparinized phosphate buffered saline (PBS) to remove red blood cells. **b** Overview of lung lobe allocation and procedure for each assay. (1) Right cranial and middle lobes were perfused post-extraction with 4% paraformaldehyde and fixed for 24 hours at room temperature. Right cranial lobes were processed for passive clarity (PACT) based clearing and immunolabelling before imaging with light sheet fluorescent microscopy (LSFM). (2) Regions of interest (ROI) imaged with higher magnification using LSFM were scanned using microCT, excised, and resin embedded prior to Serial Block Face Electron Microscopy (SBF). (3) Right caudal lobes were placed in 10% formalin and fixed for 24 hours prior to paraffin embedding and sectioning and hematoxylin and eosin staining for standard histology. (4) Left lobes were removed following cardiac perfusion, placed in 4mL PBS, homogenized and plated for CFU determination. **c** Bacterial growth and lung pathology in the lungs of C3HeB/FeJ mice exposed to a low dose aerosol infection of *M. tuberculosis* H37Rv WT vs ΔRD1 mutant over time (Day 28, 42 and 70 post-infection). (A) Colony forming units (CFU) of WT vs ΔRD1 in lung homogenates from C3HeB/FeJ mice over time following low dose aerosol infection (<100 CFU, day 1). Data points represent mean log_10_ CFU± s.e.m. of ten animals per time point (* p < 0.05; ** p < 0.01). **d** Representative histological (H&E) staining) images of lower right lobes from infected mice at day 28 (D28), day 42 (D42) and day 70 (D70). Scale bar; 1000μm (upper panels) and 50μm (lower, zoomed in panels). ZN panels correspond to parallel histological sections of the same region stained using Ziehl-Neelsen staining. **e** Percentage lung involvement at D28, D42 and D70 for uninfected animals (uninf.), ΔRD1 mutant (RD1) and H37Rv-WT indicating cell infiltration. Data points from 2 animals (uninf.) and 10 animals (RD1 and WT) per time point (* p < 0.05; ** p < 0.01; *** p < 0.001).

Bacterial growth kinetics of the E2Crimson *Mtb* WT and E2Crimson *Mtb* ΔRD1 were monitored in the lungs over time. CFU counts showed consistent growth for both the E2Crimson *Mtb* WT and E2Crimson *Mtb* ΔRD1 mutant early in the infection (day 28). However, there was significantly reduced growth observed in the mutant at day 42 and day 70 post-infection (Fig 1d). Lung histopathology analysis (Fig 1b) showed early evidence of lesions at day 28 in E2Crimson *Mtb* WT infected mice, further expanding across the lungs at later time points. By contrast, E2Crimson *Mtb* ΔRD1-infected mice showed little evidence of cell infiltration, even at later time points. Quantification of the total lung inflammation using traditional pathological scoring showed significant differences across all time points between the E2Crimson *Mtb* WT and the E2Crimson *Mtb* ΔRD1 mutant (Fig 1e). The E2Crimson *Mtb* ΔRD1 mutant had significantly less lung involvement, even at the later time points. Of the ten mice analyzed, only a small subset showed any lung involvement (Supplementary Tables S1) despite consistent growth of the strain as indicated by CFU analysis. The E2Crimson *Mtb* WT-infected mice showed a median of 1.5% lung involvement at day 28, increasing to 12% by day 42 and 18% by day 70. E2Crimson *Mtb* ΔRD1-infected mice showed almost no lung involvement throughout the infection period, as assessed by H&E (Fig 1e).

### Light sheet fluorescent microscopy (LSFM) allows the visualization of lesion development in whole lungs during *Mtb* WT infection

Following successful clearing of infected lung lobes using PACT and RIMS matching with HistoDenz™, we confirmed that the E2Crimson reporter from *Mtb* was detectable in unlabeled tissue and determined the level of resolution obtained using light sheet fluorescent microscopy (LSFM). A cleared lung lobe infected with E2Crimson *Mtb* WT (42 days post-infection) was imaged using a LaVision Ultramicroscope II Light Sheet Microscope (Miltenyi Biotec) using low magnification (0.63 X zoom) and static light sheet (12µm thickness) to generate an overview of the entire lobe, and further imaged at a higher resolution using 6.3X zoom and dynamic horizontal focusing.

We first established background autofluorescence levels, indicating a high level of tissue autofluorescence in the green channel (illumination with 488nm laser line) (Fig2a), which could be used as a marker for general airway and lung architecture without the requirement of immunolabelling. The E2Crimson reporter was visible in both the red and far-red (illumination with the 594 and 647nm laser lines respectively; data not shown). The near infra-red channel (790nm) was identified as having no autofluorescence, from either the tissue nor the E2Crimson reporter, and thus selected for further immunostaining using near infra-red fluorophores. The images showed appropriate imaging depth and enabled visualization of the entire lung anatomy (Figure 2a and Supplementary movie S1), as well as infected zones containing *Mtb* (Figure 2b). We further imaged a section of infected space using a higher magnification (6.3 X) and dynamic focus, allowing deeper visualization of infecting bacteria (Figure 2c; Supplementary movie S2). Optical sectioning allowed interrogation of different depths of the lung tissue and individual *Mtb* cells were visible in the orthogonal stack (white arrows, Figure 2C, orthogonal slices). However, the *Mtb* signal was predominantly visible as a mass, where individual spots could not be resolved accurately. Imaris cell counting was utilized to determine the total number of cells in a region and the average spot length calculated using ellipsoid axis length. On average, spot counting identified the mean length of detected spots as 4 µm (Figure 2d), with several cell spots longer that 10 µm, likely representing cell aggregates rather than single cells. This data indicates the limitation to identify and count individual *Mtb* cells with this imaging approach.

**Fig 2.**
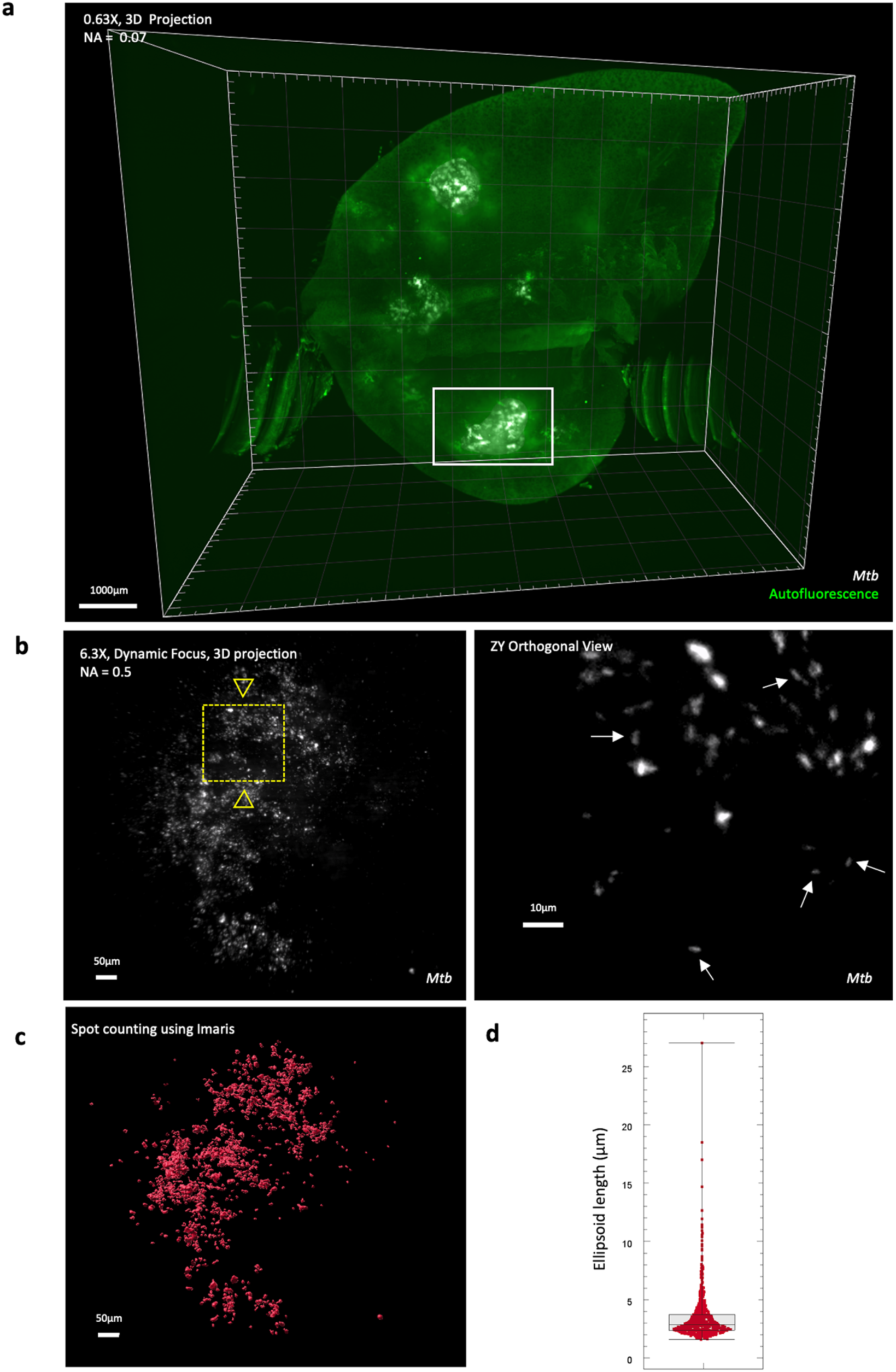
3D view of an infected (D42) lung lobe using light sheet fluorescent microscopy (LSFM). **a** Whole lobe imaging using 0.63X zoom showing the merged signal from the *M. tuberculosis* (*Mtb)* E2Crimson reporter (white) and tissue autofluorescence (green). Image was acquired with 100% sheet width (12µm thickness), 10µm z-step size and 100ms exposure time and illumination from left and right side. Scale bar = 1000µm, numerical aperture (NA) = 0.07. **b** 3D view of a region of interest (white box, panel a) imaged at higher resolution (dynamic focus with 6.3X zoom and NA of 0.5) to identify the *Mtb* E2Crimson reporter in higher detail. Image was acquired with 100% sheet width (7µm thickness) at 2µm z-step interval and 100ms exposure and illumination from left and right side. Scale bar = 50µm. The ZY orthogonal view (position 183µm) is referring to a subregion of the volume shown in fig b; showing individual *Mtb* cells (white arrows). ZY projections are enlarged roughly 5-fold relative and image gamma modified to enhance display of dim features. Scale bar = 10µm. **c** Individual cell counting using Imaris spot counting feature to identify and count individual cells from the 6.3X stack. **d** Box and whisker plot showing the average length (ellipsoid axis length in µm) of the *Mtb* cells identified during spot counting.

We next utilized PACT-based clearing and whole-lung imaging of infected animals to track infection progression over time (28-, 42- and 70-days post-infection) in 3D. To detect the majority of immune cells involved in the anti-TB response, we employed an anti-CD11b antibody (conjugated to near-IR fluorophores) to enable the identification of all myeloid cells (CD11b), coupled with the endogenous labelling of *Mtb* using the E2Crimson reporter. E2Crimson *Mtb* WT H37Rv-WT-infected C3HeB/FeJ mice showed early evidence of lesion formation at 28 days post-infection, with small foci of CD11b positive cells stemming from an upper branch (Figure 3a and Supplementary Video S3). Different from a classical single plane visualization, the orthogonal view from the 3D stack of the lungs allows the visualization of whole cell populations in defined tissue environments and *Mtb* growth (Figure 3; XY projections and Supplementary Video S3_Orthogonal), stemming from bronchioles in the regions of interest. The proliferation of *Mtb* was apparent by day 42 post-infection (Figure 3b and Supplementary Video S4) and highly evident by day 70 (Figure 3c and Supplementary Video S5). Lesion size grew substantially by day 42 and showed evidence of merging by day 70. Growth of *Mtb* at day70 showed the presence of bacteria outside of the CD11b cluster of cells, suggesting dissemination beyond the primary lesion (Figure 3c). Videos of the 3D imaged lungs of the E2Crimson *Mtb* WT H37Rv-WT-infected C3HeB/FeJ lungs can be viewed in supplementary video S3 (Day 28), S4 (Day 42) and S5 (Day 70).

**Fig 3.**
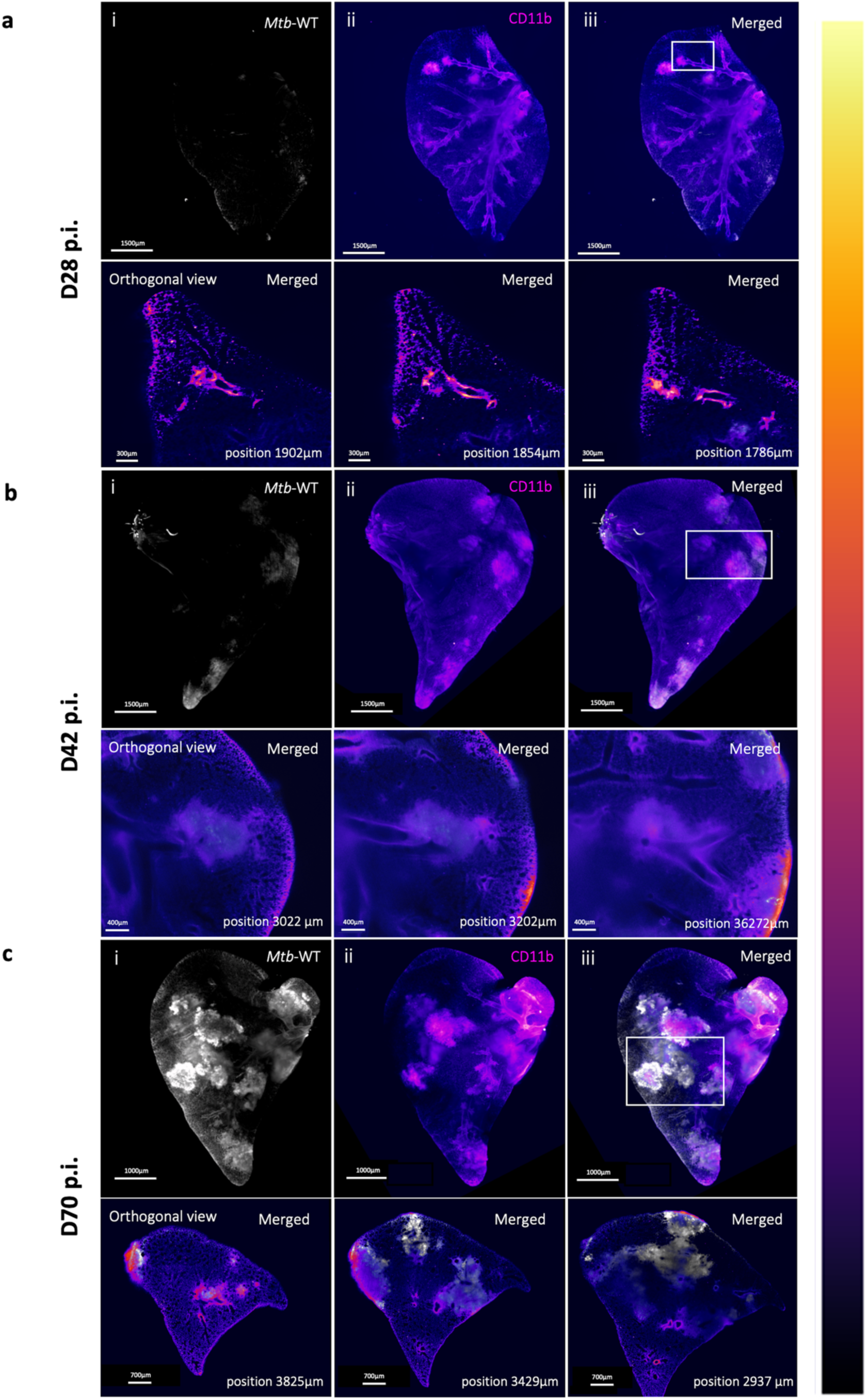
Light sheet fluorescent microscopy (LSFM) imaging of PACT-based cleared C3HeB/FeJ mouse right cranial lobe following *M. tuberculosis* E2Crimson *Mtb* WT infection over three time points; **a** D28, **b** D42 and **c** D70 (days) post-infection (p.i.). Lesion formation during disease is depicted by the accumulation of myeloid cell around infecting *Mtb* (CD11b depicted in the colour gradient lookup table ‘flame’ and E2 crimson reporter for *Mtb* H37Rv-WT shown in white). Lower panels depict ZY orthogonal positions in the lung stack highlighting specific areas of interest (white box) showing merged signal of *Mtb* and CD11b. Scale bar = 1500µm for the 3D panels and 100µm, 400µm and 700µm for the D28, D42 and D70 orthogonal sections, respectively.

We further imaged regions of interest in the E2Crimson *Mtb* WT-infected group (Figure 4a) at higher resolution using dynamic focusing (Figure 4b) and quantified the volume of a CD11b positive lesion, as well as the signal and infecting space taken up by *Mtb.* Orthogonal sections showed location of *Mtb* (Figure 4c) and using lookup table display allowed visualization of the density of CD11b surrounding the infecting cells (Figure 4d). Volume render showed lesion size grew substantially, from 5.84 x 10^7^ µm^3^ at day 28, to 2.13 x 10^8^ µm^3^ at day 42 and 1.32 x 10^9^ µm^3^ by day 70. *Mtb* occupied 0.6% of the lesion space at day 28, reaching 2% and 5.6% by day 42 and day 70, respectively with *Mtb* cells present outside of the containment regions by these time points (Figure 4 e and f). Measuring signal intensity from the E2Crimson reporter indicates the expansion of *Mtb* growth over time (Figure 4f). Videos of the total lesion render for day 70 lung lobe is provided in supplementary video S9 and individual lesion and bacterial render in S10.

**Fig 4.**
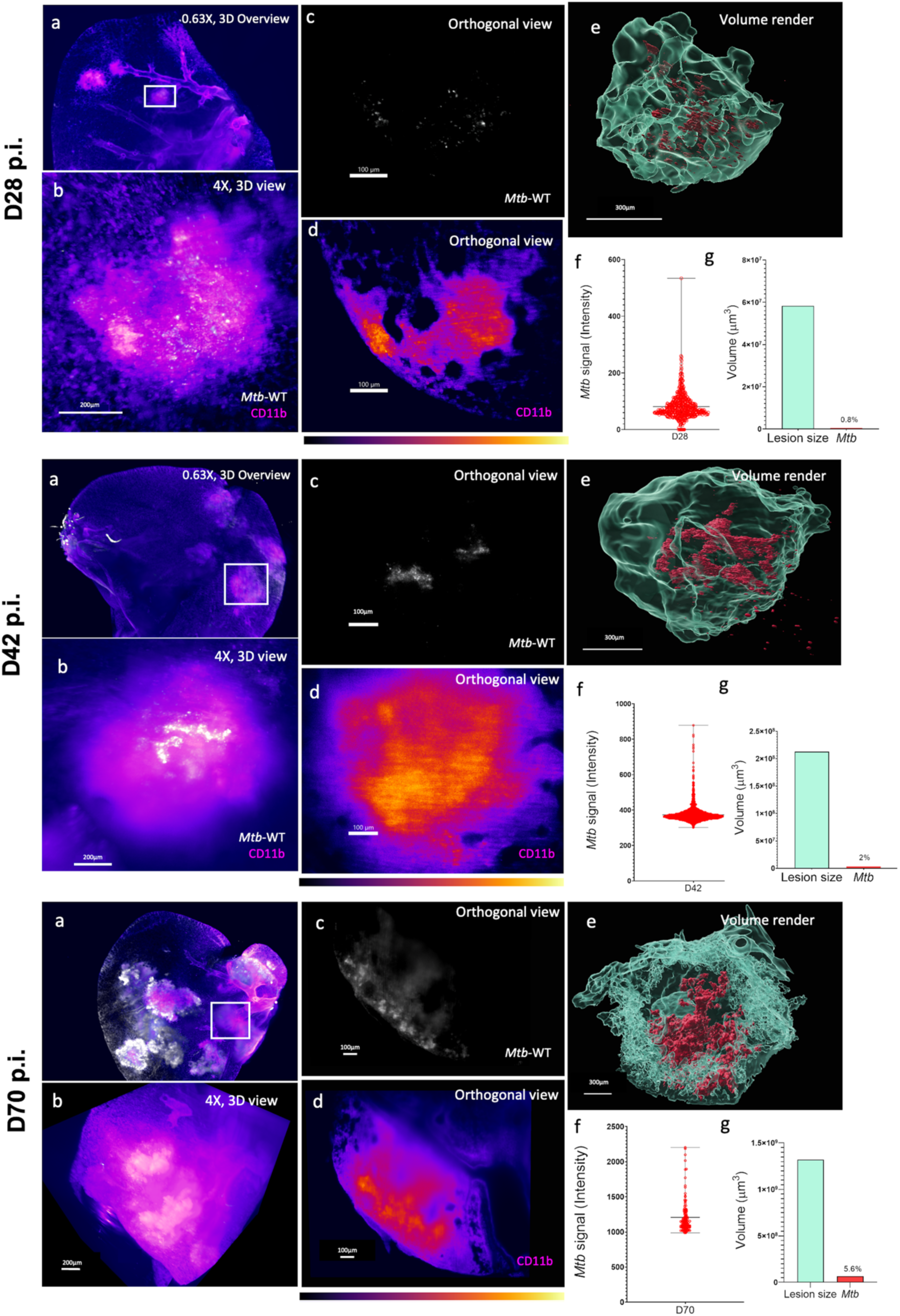
Higher resolution imaging of regions of interest from the E2Crimson *Mtb* WT-infected group of C3HeB/FeJ mice at day 28, 42 and 70 post-infection. **a** Regions of interest (ROI) identified in the larger stack under lower magnification (0.63X) are highlighter in white. **b** 3D projection of the ROI taken at higher resolution (4X, dynamic focus) showing the merged signal of *Mtb* (white) and CD11b positive cells (magenta) in lesions at day 28, 42 and 70. **c** and **d** Orthogonal view through the higher resolution stack showing the signal of *Mtb* and CD11b cells (lookup table ‘flame’ display). **e** Volume render of the lesions including render of the *Mt*-infected cells; **f** Signal intensity from the E2Crimson reporter and the volume size (µm^3^) of the lesion (cyan) and *Mtb* space (red) taken up inside the lesion.

### *Mtb* ESX-1 T7SS is required to shape lesion architecture

We next performed whole lobe imaging of E2Crimson *Mtb* ΔRD1-infected mice to compare the CD11b and *Mtb* spatio-temporal distribution pattern in whole lungs with the E2Crimson *Mtb* WT-infected mice^21^. *Mtb* utilizes specialized secretion systems to exploit host responses with the ESX-1 secretion system encoded in the RD1 region of the genome that is critical to virulence in mice. Comparison of the 3D images is shown in Fig 5 a and b and supplementary video S6 (Day 28), S7 (Day 42) and S8 (Day 70). In contrast to *Mtb* WT-infected mice-infected mice, LSFM imaging of the E2Crimson *Mtb* ΔRD1-infected mice displayed diffused signal of the *Mtb* mutant, only apparent at day 42 and day 70 post-infection. Moreover, the CD11b-enriched lesions showed disrupted, disorganized structures that were primarily present on the outer areas of the lung space with no formal structured lesion development visible (Figure 5c).

**Fig 5.**
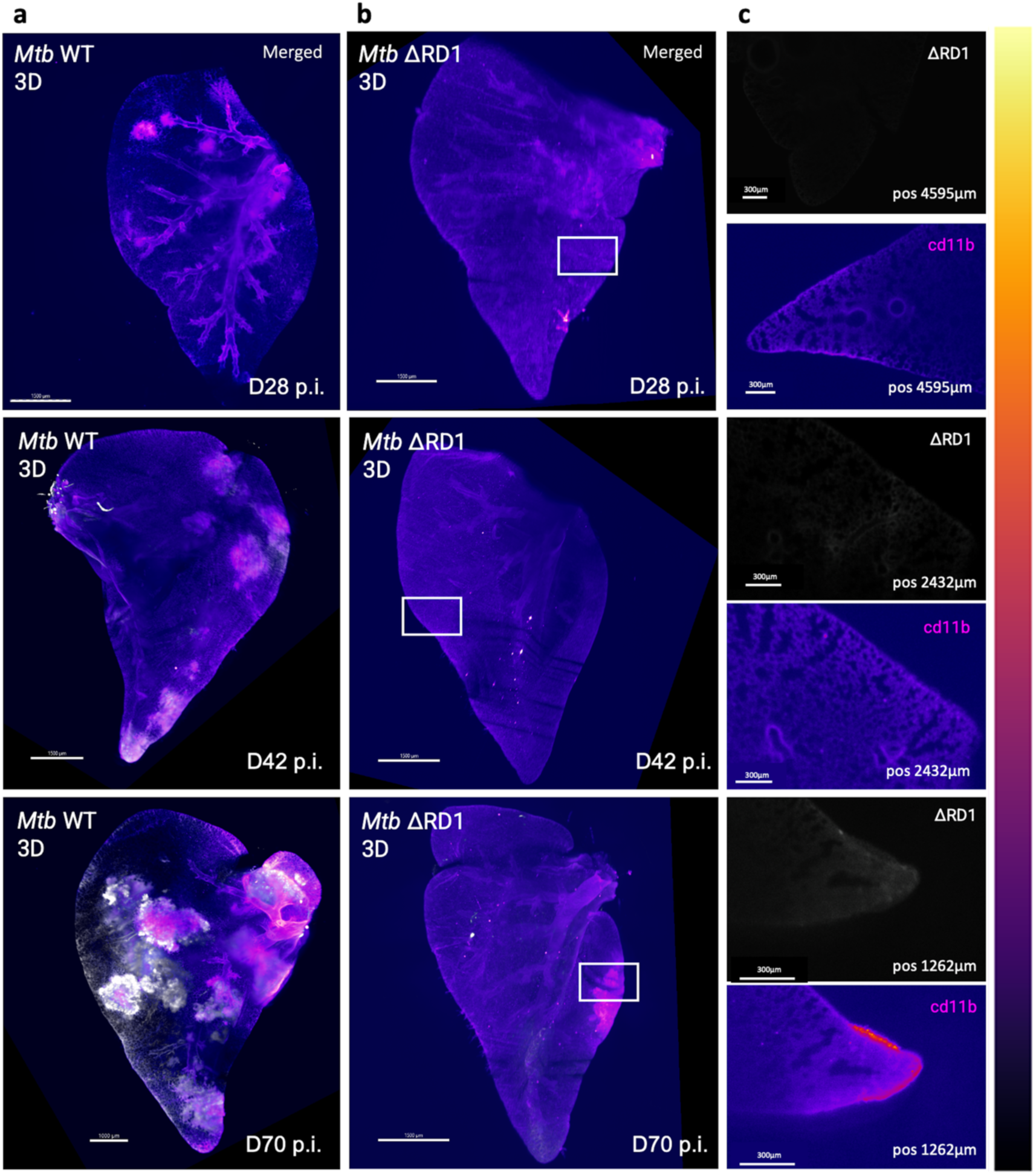
Comparison of 3D imaging of **a** E2Crimson *Mtb* WT vs **b** E2Crimson *Mtb* ΔRD1 infection over time (28, 42 and 70 days post-infection (p.i.)) in a C3HeB/FeJ mouse model depicting lesion formation during disease progression (myeloid cell recruitment depicted in magenta (CD11b) and E2Crimson reporter for *M. tuberculosis* shown in the lookup table ‘fire’). Images were acquired using identical laser and light sheet settings for both *Mtb* WT and *Mtb* ΔRD1 lung lobes. Brightness and contrast were adjusted to show visually similar intensities in the figure to account for excess intensity in the WT-infected group at later time points. **c** Zoomed orthogonal positions in the lung stack from the E2Crimson *Mtb* ΔRD1 infected lungs highlighting specific areas of interest (white box) showing signal of *Mtb* ΔRD1 and CD11b.

A quantitative analysis of lesion size and sphericity further revealed striking differences in development and progression of lesion architecture between the E2Crimson *Mtb* WT and E2Crimson *Mtb* ΔRD1 infected lungs (Figure 6). When normalized to healthy lung space, volume of infected space taken up comprised 0.4% and 0.3% of the total lung volume at day 28 for the E2Crimson *Mtb* WT and E2Crimson *Mtb* ΔRD1-infected mice, respectively (Figure 6a). Lesion volume (Figure 6b) at day 28 was similar for both *Mtb* WT and *Mtb* ΔRD1 (mean volume of lesions = 2.75 x 10^7^μm^3^ vs 2.97 x 10^7^μm^3)^ but differed substantially by day 42 (2.96 x 10^8^ μm^3^ vs 3.08 x 10^6^ μm^3^) and day 70 (3.74 x 10^8^ μm^3^ vs 8.22 x 10^7^ μm^3^). Lung involvement did not change substantially for the mutant, however in the E2Crimson *Mtb* WT infected group CD11b positive lesions comprised 7% and 10% of the total lung space at day 42 and day 70, respectively. We used sphericity measurements to quantify the spatial organization of lesions indicating spatially different lesions between the E2Crimson *Mtb* WT and E2Crimson *Mtb* ΔRD1 infected mice (Figure 6c). Altogether, using this approach we show that ESX-1 activity is required for lesion architecture in C3HeB/FeJ.

**Fig 6.**
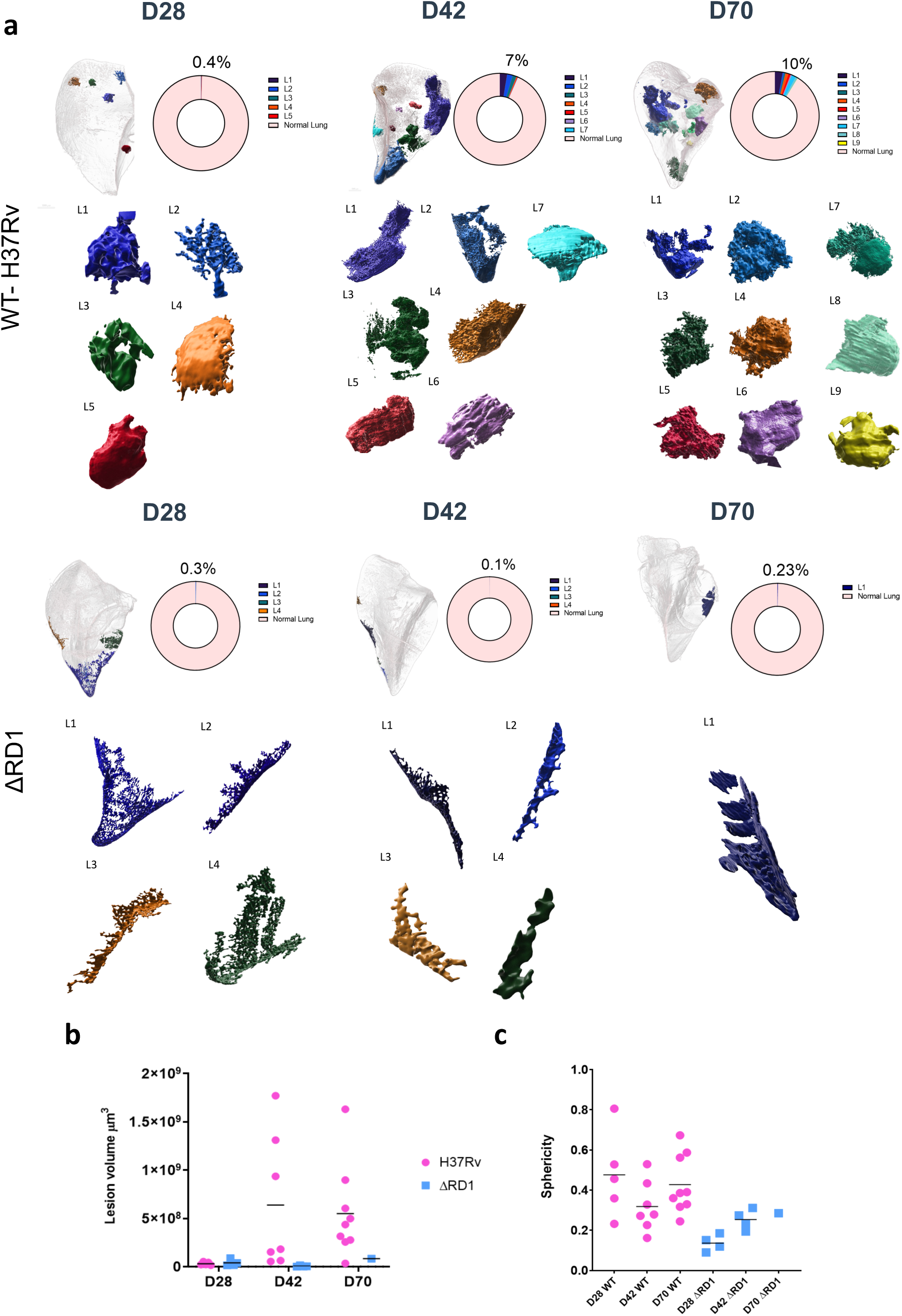
Lesion size, distribution and sphericity within C3HeB/FeJ mouse lungs infected with either *M. tuberculosis* H37Rv WT vs ΔRD1 infection over time. **a** Lesion involvement calculated as a percentage of the total lung space showing individual lesions rendered in 3D. **b** Individual lesion volumes (μm^3^) plotted over the course of infection, line bar indicated median size. **c** Sphericity of individual lesions in C3HeB/FeJ mice infected with different *M. tuberculosis* strains.

### Lesion ultrastructure through volumetric Correlative Light and Electron Microscopy (vCLEM) shows that infection spreads from an initial macrophage population

PACT clearing allowed for large scale characterization of entire lung lobes and molecular identity through immunolabelling and LSFM. Given the abundance of E2Crimson signal at defined regions, we investigated whether there was a structural pattern to the observed clustering of the E2-crimson reporter at the ultrastructural level. Here, we opted to employ volumetric Correlative Light and Electron Microscopy (vCLEM) to assess lesion ultrastructure in 3D at the subcellular level. This required the development of a gentle electron microscopy staining protocol (ClearEM), capable of preserving ultrastructural detail in cleared lung tissue. To confirm whether *Mtb* could still be observed post clearing, high-resolution SEM was conducted on 100nm sections from uncleared (Fig 7a, i) and cleared *Mtb* (Fig 7a, ii) -infected mouse lung samples. After confirming the presence of bacteria within cleared tissue samples, we proceeded with our vCLEM workflow, outlined in Fig 7b. Following LSFM (Fig 7b, ii), the lung lobes were embedded in resin (Fig 7b, iv) and orientated using the LSFM data to further image using microCT (Fig 7b, v) to determine block trimming for Serial-Block Face SEM. Inherent morphological features were used as fiducial markers for multi-modal image registration and alignment. Airways and blood vessels could be used as landmarks to align the fluorescent data with the microCT data sets (Fig 7b, vi) to allow for a final alignment overlay (Figure 7c), as shown before with similar methods ^22^.

**Fig 7.**
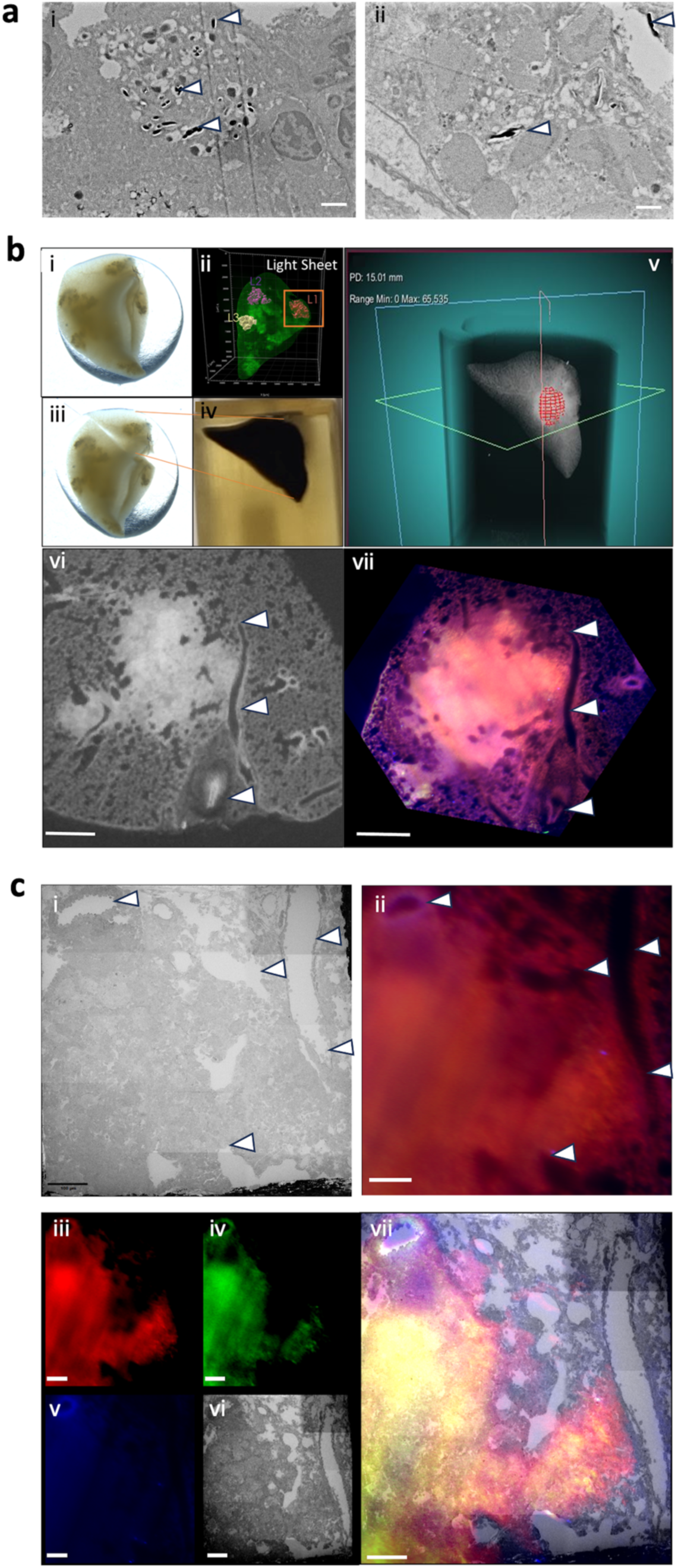
Volumetric correlative light and electron microscopy (vCLEM) to interrogate PACT-cleared tissue for ultrastructural content imaging. **a** Scanning Electron Microscopy micrographs of 100nm sections captured on silicon nano-wafers. **i** Uncleared tissue prepared with a conventional mega-metal staining protocol. White arrows indicate *Mtb* bacteria within distinct vacuoles, indicative of engulfment by macrophages. **ii** Cleared tissue prepared with ClearEM staining protocol. White arrows indicate presence of *Mtb* within similar vacuolar structures seen in uncleared tissue. Scale bar 1μm. **b** Overview of region identification workflow. **i** Entire lung lobe with **ii** corresponding light sheet render of each lesion of interest. **iii** Lung lobes were cut according to the localisation of each lesion and **iv** embedded in resin. Care was taken to leave enough healthy surrounding tissue until fine block trimming could occur post resin embedding. **v** Prior to block trimming, micro-CT scanning was performed on embedded tissue samples to assess the orientation of granulomas compared to initial light sheet microscopy. Image registration between the micro-CT and light sheet microscopy was conducted to aid in block trimming. **vi** Light sheet data was transformed to micro-CT data using BigWarp (FIJI), to geometrically scale the light sheet data to the same as that in the resin block. An affine transformation was applied to the stack, with special care taken not to induce excessive deformation of the original image. The newly transformed light sheet stack was then registered to the captured SBF-SEM dataset, again using inherent morphological landmarks in order to identify appropriate areas for segmentation. Scale bar 500μm. **c** Use of morphological landmarks to overlay fluorescent and serial block face electron microscope datasets. Fine registration of **i** SBF-SEM and **ii** Light Sheet images using inherent morphological landmarks indicated by white arrows. Final channel overlays post autofluorescence subtraction is indicated in **iii, iv** and **v** and the final overlay with **vi** shown in vii. Scale bar 100μm.

Tracking the cell population corresponding to the localisation of E2Crimson fluorescent signal revealed distinct clusters of cells that are less electron dense than the surrounding tissue (Fig 8). Furthermore, these domains appear to propagate from surrounding airways, indicating that infection spread from the initial macrophage population attracted to the site of infection closest to the airways. However, these domains are distinct in that a single lesion comprises of multiple macrophage aggregates, rather than an individual amalgamation of cells, as previously thought. These results provide a unique insight into granuloma formation, and suggest that lesions are comprised of distinct macrophage populations within a single granuloma.

**Fig 8.**
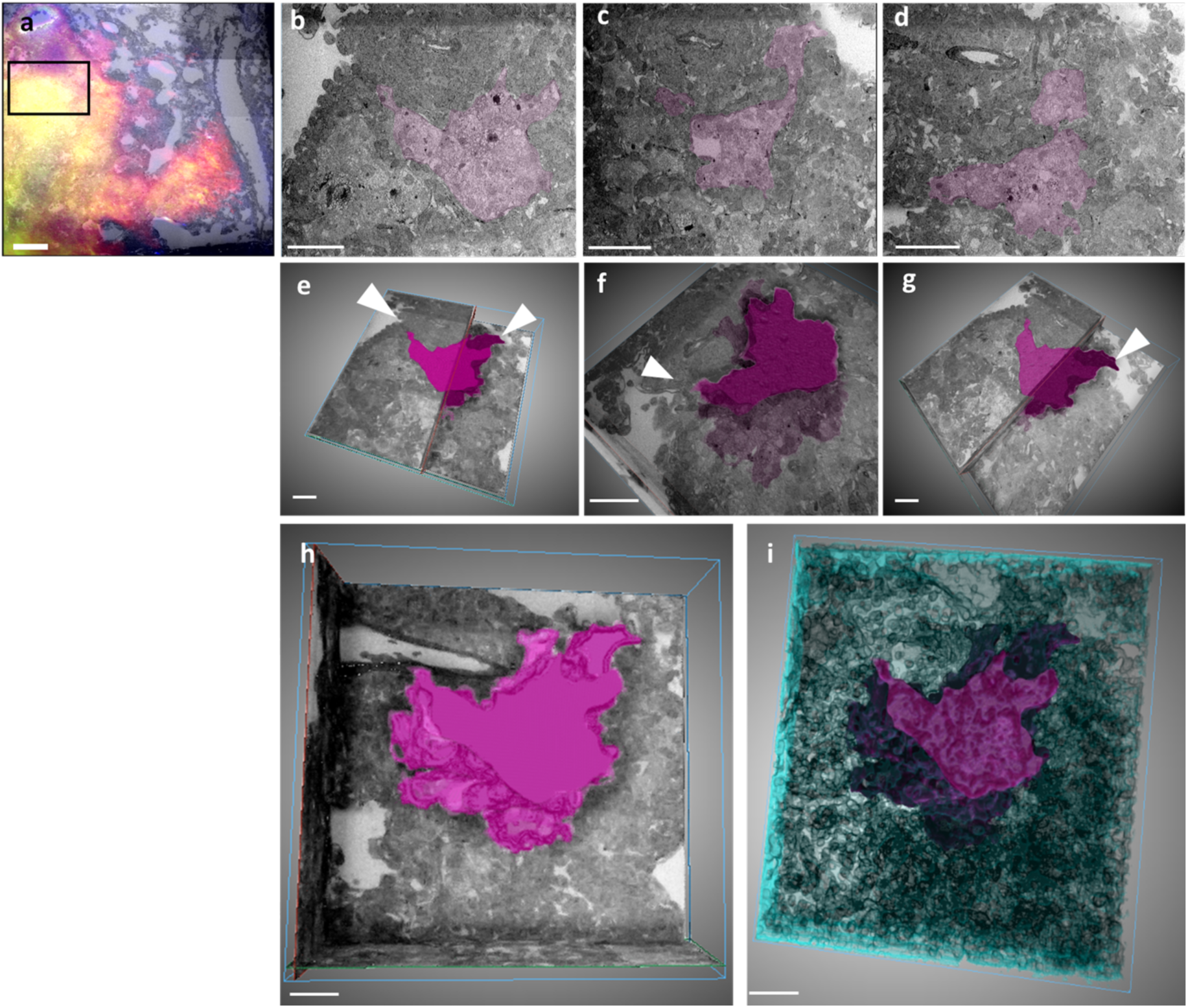
Ultrastructural analysis and tracking of cell populations in infected lesions using serial block face electron microscopy. Tracking of individual infected macrophage populations across multiple z-planes at **a** z=0μm, **b** z=300μm and **c** z=600μm. **e-g** represent total volume render of infected macrophage population within a selected lesion, with branch points from airways indicated by white arrows. **h** Render of infected population only with **i** showing the volume of surrounding structures in blue. Scale bars = 100μm.

## Discussion

Understanding the spatial architecture of granulomas is critical to our understanding of TB pathogenesis and disease progression. Here, we developed a method that allows the investigation of the spatial localization of bacteria in lesions in a mouse model of TB. Using this method, we found critical architectural differences in lesion development and lung involvement between wild type *M. tuberculosis* and an ESX-1 deletion (Mtb ΔRD1) mutant strain.

We have employed PACT clearing and LSFM to investigate lesion development in 3D for the first time in C3HeB/FeJ mice under different *Mtb* virulence conditions. LSFM provided accurate spatial anatomical information when compared to standard histology and unveiled differences in the architectural structure of lesions highlighting differences in *Mtb* type 7 secretion system ESX-1 mediated pathogenesis. Quantification of total lesion size and shape, as well as the spatial distribution in the total lung space provides a more contextual view of disease progression in a murine model. This study builds on previous studies that have utilized tissue clarification processes to investigate *Mtb* infection in whole lungs ^16,17^. The method developed here provides spatiotemporal resolution and the ability to track infection and the associated immune response in an anatomical context throughout the whole organ.

This approach provides a unique opportunity to look at host-pathogen interaction in different anatomical areas of the same lung at the same time. Combining immunolabelling, LSFM and volumetric EM modalities allows a multi-modal vantage point of infection progression and an unbiased view of pathophysiology, e.g. at the single lesion level.

Although we were limited by the available fluorescent channels, LSFM imaging allows faster and deeper imaging than can be achieved with standard confocal microscopy and without the requirement of sectioning of the tissue. This, however, comes at a resolution cost since the long working distance of the objective lens limits spatial resolution that can be achieved for an entire lung lobe. Nevertheless, we showed that higher resolution imaging of regions of interest can be achieved through higher magnification (6.3X zoom; 0.5 NA), together with dynamic focus mode and thin light sheet, although this comes at a cost in acquisition time. A higher NA allowed the visualization of individual *Mtb* cells in the orthogonal view of specific lesions although, most of the *Mtb* signal is difficult to resolve as single cells, specifically at later time points. Signal intensity can be utilized as a general marker of proliferation and with the advancements of metabolic probes for *Mtb*^23^, could be utilized in future studies to provide increased information on the state of the pathogen in different infecting zones.

Visualizing the dynamics of lesions development under different virulence conditions allowed unprecedented view of the 3D organization of lesions in C3HeB/FeJ mice. We used an ESX-1 deletion mutant to highlight the key differences between different strains during early and late lesion formation. The ESX-1 secretion system is well characterized and has been shown to play a critical role in early innate responses during *Mtb* infection^21,24,25^. ESX-1 is required for phagosome membrane damage, arresting phagosomal acidification and allowing access to the host cytosol where free replication can resume. The interaction of *Mtb* with the host has a clear impact on the outcome of infection and overall pathophysiology. Differences between early and late time points post-infection in ESX-1 mediated autophagic processes suggest that these processes are critical in sustaining the survival of *Mtb* long-term in the host. Interestingly, although CFU counts showed consistent growth of the ESX-1 mutant in homogenized tissue, standard histology of thin sections showed little to no evidence of cell infiltration. By comparison, LSFM imaging of ΔRD1-infected lung lobes unveiled the altered growth pattern of this strain in C3HeB/FeJ lungs as diffuse and sparsely organized cd11b recruitment to sites of infection. Most lesions were present on the outer areas of the lungs and did not display any formal organization. In contrast, WT-infected mice showed well-organized lung lesion structure growing substantially throughout the infection period, invading a high percentage of the total lung volume. The WT *Mtb* strain was clearly visible at all time points and increased substantially by day 70 where it was clearly visible outside of containment areas. The development of large lung lesions in C3HeB/FeJ mice when infected with a low dose aerosol WT strain in this study is characteristic of what has been observed in other studies^20,26^. Here, we showed that early infection with virulent *Mtb* begins with small foci of infection stemming from the upper airways, which eventually disseminates to the lower areas of the lung. This is in line with studies that have shown that *Mtb* virulence proteins drives tissue destruction early on and allows infected macrophages to depart the primary granuloma and disseminate the infection^27^.

One of the main limitations of this study is the laborious and time-consuming procedures required to prepare samples, which is impractical to assess large number of animals at one time. PACT clearing and LSFM imaging should thus be utilized for exploratory studies that seek spatial understanding of lesion distribution in the total lung space. In addition, immunolabelling takes an extremely long time to reach the tissue and requires large amounts of antibody. Although methods to speed up this process have been proposed such as ACT-PRESTO^28^, we did not find this to work in our experimental setup (data not shown). We propose that transgenic mice that express fluorescent reporters in cells of interest would be a more practical method to visualize and circumvent lengthy antibody incubation times.

LSFM enabled whole lobe imaging, yet the level of resolution at the cellular level is compromised. We wanted to investigate whether volumetric electron microscopy could be used to further reveal ultrastructural detail in PACT-cleared tissue which has not previously been described. Although limited to a single lesion, we were able to show that clarified tissue can be further utilized for ultrastructural imaging, albeit with a loss of membrane integrity. To our knowledge, this is the first time PACT clearing and SBF has been combined in a single study.

Altogether, we have shown that PACT clearing and LSFM imaging can provide unbiased and a contextually relevant view into lesion development in a physiologically relevant murine model of TB. LSFM can comprehensively describe all regions of infection and allows identification of important features of pathophysiology under different virulent conditions. This opens novel ways of interrogating new drugs and host directed therapies, whilst gaining insight into the host immune response.

## Materials and methods

### Mice

Research protocols adhered to guidelines set out in the South African National Standard 10386 (2021), were approved by Stellenbosch University Research Ethics Committee Animal Care and Use and all procedures were conducted according to the Veterinary and Paraveterinary Professions Act (Act 19 of 1982) and the Animal Diseases Act (Act 35 of 1984). Specific pathogen-free C3HeB/FeJ mice were purchased from the Jackson Laboratory (Bar Harbor, ME). Female mice (between 8-10 weeks of age) were housed at 5 mice per cage in individually ventilated microisolator cages under biosafety level III containment under sterile condition with sterile bedding, water and standard mouse chow. Infected mice were monitored daily for welfare and weighed every second week.

### Bacterial strains

The laboratory strain of *M. tuberculosis* (H37Rv) and an ESX-1 deletion mutant (ΔRD1) both transformed with a PTEC-19 plasmid containing an E2-crimson fluorescent reporter and hygromycin resistance cassette were used for aerosol infections. Bacteria were grown in Middlebrook 7H9 medium containing 0.05% tween and supplemented with 10% OADC (oleic acid, albumin, dextrose, and catalase; BD Biosciences) and 50ug/uL Hygromycin B (Gibco, ™) in vented flasks and kept stationary at 37°C until mid-exponential phase was reached (OD600 nm 0.6–1.0) and enumerated by colony counting on 7H11 agar plates (described below). Culture stocks were stored at -80°C until use.

### Aerosol Infection

Eight- to ten-week-old mice (40/infecting strain) were infected utilizing an inhalation exposure system (Glas-Col Inc., Terre Haute, IN). Bacterial strains from frozen stocks (H37Rv-WT and ΔRD1 mutant) were thawed and suspended repetitively using a Luer lock syringe attached to a 26 Ga hypodermic needle to obtain a single cell suspension. Infection inocula were prepared to a final concentration of 2 x 10^6^ CFU/mL in autoclaved deionized water and 5mL of this inoculum was placed in the Glas-Col nebulizer. The infection procedures included 10 compressed air and 60 main (negative) standard cubic feet per hour (SCFH); 15 min preheat time; 30 min nebulizing time; 45 min cloud decay time and 15 min decontamination time.

Enumeration of the bacterial inoculum for both strains was determined by CFU counts on 7H11 plates and the actual bacterial load delivered to the animals was confirmed from ten mice/strain one day post-aerosol challenge in total lung homogenate.

### Necropsy and organ harvesting

Following euthanasia using inhalation anaesthetic overdose, mice were individually transcardially perfused using 10mL heparinized saline solution until lungs inflated and turned white, indicating red blood cell removal. Whole left lung lobes were removed and placed in 5mL sterile PBS for enumeration of bacterial load. Right cranial and middle lobes were placed in 4% PBS-buffered paraformaldehyde and caudal lobes were placed in 10% formalin for FFPE embedding and histology.

### Enumeration of bacterial load of lungs

Lung lobes were disrupted in 5mL sterile PBS containing 1.4mm ceramic (zirconium oxide) beads (Bertin Instruments) using a tissue homogenizer (Precellys Evolution, Bertin Instruments), serially diluted, and plated onto Middlebrook 7H11 agar plates supplemented with OADC. Colonies were counted after 21 days incubation at 37°C. Bacterial counts were log_10_ transformed and expressed as mean log10 CFU ± standard error of the mean for each group.

### Pathology of lungs

Right caudal lobes were embedded in paraffin and cut to 5µm thickness using a microtome before mounting onto glass slides. Sections were deparaffinized and stained with haemotoxylin and eosin for standard pathological assessment. Slides were scanned on a Grundium Ocus® 40 scanner with a UplanXApo 20 X objective (NA 0.75; resolution 10μm/pix; image sensor: 12Mpixel). Analysis of histology slides from each timepoint was done using QuPath v0.3.2 open-source software (https://doi.org/10.1038/s41598-017-17204-5). This software was used to quantify the percentage lung involvement of lesions following *Mtb* infection vs and ΔRD1 mutant. For each image used in the analysis, the magic brush tool was used to annotate the respective section of the image. A separate copy of the image was made whereby only, if present, lung lesions were annotated used the same method. Specific adjustments were made for each annotation for each image to match the anatomy of each section and for consistency. The QuPath cell detection function was used to quantify and mark cells within an annotated area.

For each annotation, the number of detections and area were exported and analysed. Analysis for percentage lung involvement was completed by dividing the number of cell detections in lesions by the overall number of cell detections and then multiplying by 100. One-way ANOVA was performed with a Kruskal-Wallis post-test. The graph represents the results as mean ± SEM, ****. P = < 0.0001.

### Tissue Clarification

PFA-fixed samples were transferred to freshly prepared hydrogel monomer solution (A4P4, 4% acrylamide (Sigma-Aldrich Inc, USA) in 0.1M PBS) containing 0.25% photoinitiator VA-044 (Wako chemicals USA, Inc, USA) and incubated at 4°C for 48 hours to allow permeation into the tissues. Samples were degassed for 20 minutes by bubbling nitrogen gas in the medium and placed at 37°C for 4 hours for embedding. Following polymerization of the monomer, lungs were carefully removed from the hydrogel and rinsed in 0.1M PBS and transferred to clearing solution (8% sodium dodecyl sulfate, SDS, Sigma-Aldrich Inc, USA) in 0.1M PBS containing 50mM sodium sulfite (Sigma-Aldrich Inc, USA), pH 8.5. Samples were incubated in the dark at 37°C with gentle shaking (100rpm) and clearing solution was replaced daily. Once optical transparency was achieved (approximately 4 days), SDS was washed out with sequential wash steps in wash buffer (0.1% Triton-X100 in 0.1M PBS containing 50mM sodium sulfite, ph 8.5) by diluting the clearing solution 1:1 with pre-warmed wash buffer three times daily for 2 days, gradually reducing the temperature until room temperature was reached. Samples were stored in 0.1M PBS containing 0.01% sodium azide before immunostaining and imaging.

### Immunostaining

Lung lobes were washed twice for 20 min using 0.1M PBS and transferred into 10mL/lobe blocking buffer (3% BSA in 0.1M PBS containing 0.1% Triton X-100 and 0.01% sodium azide) and incubated at 37°C with gentle shaking for 2 days in the dark. Lung lobes were stained for a total period of 5 days, progressively increasing antibody concentration every day, reaching final concentration of 1:50 by day 4. This was done to prevent antibody accumulation on the outside of the lungs which was observed during optimization. A total volume of 3mL primary antibody (anti-cd11b, Invitrogen, Cat no PA5-79532) in blocking buffer was used to stain each lung lobe at 37°C with gentle shaking for 4 days in the dark. Following immunostaining, lung lobes were washed for 3 days in 50mL 0.1M PBS containing 0.1% Triton X-100 at room temperature, changing the wash solution twice daily. Lung lobes were then transferred to secondary antibody solution (1:50 Alexa Fluor 790 AffiniPure Fab Fragment Goat anti-rabbit, Jackson Immunoresearch Laboratories, Cat no: 111-657-008), in blocking buffer and incubated for a further 2 days at 37°C with gentle shaking for in the dark. A further 2 days of washing in 0.1M PBS containing 0.1% Triton X-100 was conducted to wash off unbound secondary antibody. All blocking buffer solutions were filter sterilized using a 0.22µm filter (Sigma-Aldrich).

### RIMS matching

Refractive index (RI) matching was done by immersing the stained lung lobes in home-made RIMS at an RI of 1.45. Briefly, 143.78g of HistoDenz™ (Sigma-Aldrich) was dissolved in 0.02M PB buffer containing 0.1% Tween-20 and 0.01% sodium azide, pH was adjusted to 7.5. A refractometer was used to confirm the RI of the solution. Lung lobes were incubated in excess RIMS solution (5mL) in the dark until completely transparent. The same solution was used to fill the imaging chamber during imaging.

### Light Sheet Fluorescent Microscopy and data acquisition

Lung lobes were mounted directly onto platforms with either screws or spikes to fix the sample in place, ensuring the sample was stable throughout acquisition. Sample orientation was carefully chosen to ensure that propagation length of the excitation light is a short as possible, whilst maintaining the full ROI does not exceed the working distance of the objective. The chamber was filled with 130mL of the RIMS matching solution. Single plane illuminated (light-sheet) image stacks of cleared lung lobes were acquired using a Miltenyi-LaVision Biotec Ultramicroscope II Light Sheet system (Miltenyi Biotec GmbH, Bergisch Gladbach, Germany) equipped with Olympus MVX-10 Illumination microscope body and MV PLAPO 2XC (Olympus, Tokyo, Japan) objective. Samples were illuminated by three light sheets from left and right side. Whole lung lobes were acquired at 0.63X zoom with an exposure time of 100ms, z-stack intervals of 6µm and light sheet thickness of 12µm. The *Mtb* E2-crimson reporter was imaged with 640nm laser line and 680/30 emission filter with laser power at 90% and cd11b signal was imaged using 784nm laser line and 835/70 emission filter with laser power of 100%. Where autofluorescent signal was required, green signal was imaged using the 488nm laser (excitation) and 525/50 emission filter with 20% laser power. Higher resolution images were taken on areas of interest using either the 4X or 6.3X zoom depending on where the lesion was situated and the working distance available. Dynamic focus allowed capturing of high-resolution images across a large area. Instead of keeping the light sheet in one location and moving the sample to tile the image, dynamic focusing keeps the sample stationary and adjusts the horizontal focus position of the sheet throughout it. Multiple images are acquired and then blended to ensure maximal Z resolution across the entire field of view. Dynamic horizontal focusing mode was employed with blending mode set to the center of the image with light sheet thickness adjusted to 3.8μm and a step size of 0.5μm. ImSpector Pro software was used for imaging and IMARIS software v9.7.2 (Bitplane AG, CA, USA) was employed to analyse LSFM images. Image stacks were converted to native ims. files using file converter tool and used to generate 3D video files. The surface creation tool was used to create a region of interest to set threshold signals after which the entire image was processed. In the surface creation wizard, the source channel for either the E2-crimson reporter or AF790 channel was selected. Smoothing surface detail was set to 7µm. Maximum intensity was used to identify the cell of interest in each lobe, number of voxels was kept at 1mg and gamma as 1.00. Adjustments to surface shape and detection were made manually to reduce extraneous noise, particularly where the signal was oversaturated in the later time points. Spot counting was used using the Imaris spot component, segmenting region of interest for analysis and using “different spot size” feature to assign different sizes based on the adjusted signals. Estimated XY diameter was assigned as 5µm and background subtraction (Gaussian) was enabled. Model PSF-elongation along z axis was enabled to created elliptical shaped spots. Threshold was adjusted manually to have as many single spots as possible per identification. Individual lesion size and sphericity was calculated using the surface object detection and 3D volume rendering tool with blend mode with manual thresholding of intensity based on individual signal in each lobe. Threshold signals set as follows: H37Rv WT D28 = max 2102, min 511; D42 = max 2160, min 511; D70 = max 1241, min 241 and RD1 mutant: D28= max 4849, min 249, D42 = max 6697, min 249, D70 = max 2551, min 249. Zoomed lesions were analyzed similarly, with threshold settings as follows: D28 = max 534, min 0; D42 = max 879, min 302, D70 = max 7969, min 987.

### Volumetric Correlative Light and Electron Microscopy (vCLEM)

#### Sample Preparation for Serial Block-Face Electron Microscopy

Following light sheet microscopy, lung lobes were cut according to the localisation of each lesion. Care was taken to leave enough healthy surrounding tissue until fine block trimming could occur post resin embedding. Thereafter, samples were fixed for 24 hours at 4 °C with a mixture of 2.5 % glutaraldehyde and 4 % formaldehyde in 0.1 M phosphate buffer. Thereafter, incubation with 2 % reduced osmium tetroxide (OsO_4_) (mixture of 4 % OsO_4_ and 3 % potassium ferricyanide, 1:1) was conducted for 60 minutes on ice, followed by 20-minute incubation with thiocarbohydrazide (TCH) at room temperature, 30 minutes incubation with aqueous osmium tetroxide at room temperature and overnight incubation with 1 % uranyl acetate at 4°C. Between each incubation step samples, samples were washed three times with distilled water (dH_2_O).

After overnight incubation, dehydration in a graded ethanol series of increasing concentrations was conducted for 10 minutes each on ice (20 %, 50 %, 70 %, 90 %, 100 %), followed by two rounds of 100 % ethanol dehydration at room temperature. Resin infiltration was performed in a series of 2-hour incubation steps at room temperature, using increasing concentrations of EPON resin diluted in Acetonitrile (25 %, 50 % and 75 %). A final infiltration step with 100 % EPON resin was performed overnight on a rotator, followed by incubation with fresh resin for 4 hours the next day. Lastly, samples were transferred to embedding mounts and placed in an oven at 60 °C for 2 days to allow for appropriate polymerisation and hardening of resin.

#### Micro Computed Tomography (CT) Scanning

Prior to block trimming, micro-CT scanning was performed on embedded tissue samples to assess the orientation of granulomas compared to initial light sheet microscopy.

#### Region finding and block trimming

Resin blocks were then trimmed down coarsely by hand, followed by fine trimming of an initial region of interest (ROI) spanning 2 x 2 x 1 mm using a glass knife and Leica UC7 ultramicrotome (Leica Microsystems, Austria). The trimmed block was then mounted on an ultramicrotome stub using conductive silver epoxy (SPI, S-05000-AB). To facilitate hardening of the silver epoxy, samples were transferred to an oven for 2 hours at 60 °C. Excess epoxy was trimmed away, and fine cutting of the resin block continued with the ultramicrotome, this time using a 45° diamond knife (Diatome, USA). When the block face reached an approximate area of 0.7 x 0.7 mm, 100 nm thin sections were captured on silicon nano-wafers to asses tissue ultrastructure through SEM (ThermoFisher Apreo SEM). Once the region of interest was confirmed to be a granuloma, with corresponding morphological landmarks to that of the micro-CT data, the block-face was sputter coated with 50 nm gold/palladium (Denton Desk V, USA) followed by another round of ultramicrotome sectioning (80 nm thin sections) to expose the sample block-face.

The sample stub was placed into the ultramicrotome attachment of the Apreo Volumescope (ThermoFisher, Netherlands) and eucentric calibration of the diamond knife to the block-face was conducted. Image acquisition was conducted at an adjusted chamber pressure of 0.5 mbar to compensate for excessive charging of the resin. Under a lowered chamber vacuum pressure of 0.5 mbar, the VolumeScope Dual Back Scatter (VS DBS) detector was implemented, using a voltage of 2.00 kV and probe current of 0.20 nA. Region finding and beam energy alignment were conducted using Xt Microscopy and Maps 3.9 software (ThermoScientific). A z-width of 100 nm was deemed appropriate as cutting thickness. Serial block-face data was acquired for a total of 2000 slices using the VS-DBS detector at a pixel resolution of 3840 x 2160 for each tile with a scan speed of 2 µs.

#### Image Processing

After acquisition, tiled z-stacks were stitched using the MIST plugin in FIJI(ImageJ) followed by SIFT linear alignment of z-stacks in FIJI(ImageJ). Image processing involved applying a 2D gaussian blur (Sigma=2), recursive exponential filter (3×3×3 pixels) and a final 3D denoising filter (5×5×5 pixels) in Amria (2019.3).

Image registration between the 3 modalities involved transforming light sheet data to micro-CT image stacks using BigWarp (FIJI), to geometrically scale the light sheet data to the same as that in the resin block. An affine transformation was applied to the stack, with special care taken not to induce excessive deformation of the original image. Given that a subsequent transformation would be necessary for alignment to the SBF-SEM data, a similar transformation was deemed unfit. The newly transformed light sheet stack was then registered to the captured SBF-SEM dataset, again using inherent morphological landmarks in order to identify appropriate areas for segmentation.

Once identified, manual segmentation was conducted on regions of interest using ORS Dragonfly (2022.1), with all subsequent renders and image panels also created in the same software.

## Statistical analysis

All statistical analysis was performed using GraphPad Prism version 7. Specific tests employed are described in each figure legend.

## Data availability

Due to their large size, the raw imaging data that supports the findings of this study are directly available from the corresponding author upon reasonable request. Derived data have been compiled in the Source Data file provided with this paper.

## Acknowledgements

This work was supported by the Crick African Network which receives its funding from the UK’s Global Challenges Research Fund (MR/P028071/1), and by the Francis Crick Institute which receives its core funding from Cancer Research UK (CC2081), the UK Medical Research Council (CC2081), and the Wellcome Trust (CC2081) to MGG. For the purpose of Open Access, the author has applied a CC by public copyright licence to any Author Accepted Manuscript version arising from this submission. The authors are grateful to Dr. Lucy Collinson and Dr. Marie-Charlotte Domart at the Crick EM STP for their expertise and assistance in preparing and conducting the micro-CT work, and Profs. Anne Lenaerts and Greg Robertson for their expert contribution to guiding the application of the C3HeB/Fej mouse model and Dr Verena Wimmer from the Centre for Dynamic Imaging at WEHI for her expert guidance in tissue clarification. The authors would also like to thank the Stellenbosch Animal Facility, especially Dr Este Spies, Mr Noel Markgraaff, Mr Romano Markgraaf, and Mr Abraham Slinger for their invaluable assistance with the mouse work, as well as Mr Reggie Williams and Mr Jeffrey Pietersen for their contribution to the histopathology work.

## Notes

### Competing Interest Statement

The authors have declared no competing interest.

